# Genotypic Characterization of Virulence and Antibiotic–Heavy Metal Resistance in Enterococcus Strains from Vladivostok’s Water Bodies

**DOI:** 10.1101/2025.05.26.656074

**Authors:** Jun Hyung Kim, Mayuri Hujuri, Svetlana Sergeevna Uskova, Alina Viktorovna Martynova

## Abstract

1.

**Background:** *Enterococcus* species are Gram-positive, facultative anaerobic bacteria commonly found in soil, water, and the gastrointestinal tracts of humans and animals. Their ability to acquire antibiotic resistance has made them significant opportunistic pathogens in both clinical and environmental settings. The presence of antibiotic-resistant *Enterococcus* strains in aquatic environments reflects contamination from human activities and poses a potential risk of horizontal gene transfer to other bacterial species.

**Aims:** This study aims to assess the antibiotic and heavy metal resistance patterns of Enterococcus strains isolated from water bodies in Vladivostok and evaluate their potential environmental risks.

**Study design:** Experimental study.

**Methods:** Water samples were collected from the Vtoraya Rechka River and Zolotoy Rog Bay in Vladivostok. 30 *enterococcus* strains were identified by polymerase chain reaction (PCR) technique. The presence of antibiotic and heavy metal resistance genes was determined using specific primers.

**Results:** Genetic analysis of the isolated Enterococcus strains revealed that 23% of strains carried the *tetL* gene, 33% harbored the *ermB* gene, and the heavy metal resistance genes *tcrB* and *cadA* were detected in 40% and 27% of strains, respectively. Notably, the *ermB* and *tcrB* genes co-occurred in 13.3% of strains, while none of the *tetL*-positive strains exhibited simultaneous presence of *ermB, tcrB*, and *cadA*.

**Conclusion:** *Enterococcus* strains isolated from Vladivostok’s water bodies exhibit resistance to multiple antibiotics and heavy metals, with a notable co-occurrence of copper and antibiotic resistance genes. These findings contribute to understanding microbial contamination in aquatic environments and its public health implications, providing valuable insights for future research in clinical and environmental microbiology.

**IMPORTANCE:** The dissemination of antibiotic and heavy metal resistance genes in environmental reservoirs poses a serious public health threat, as these genes may be transferred to clinically relevant pathogens. *Enterococcus* species, widely present in aquatic ecosystems, serve as both reservoirs and indicators of such resistance. This study identifies patterns of co-occurrence among resistance genes and highlights a potential genetic linkage between macrolide (*ermB*) and copper (*tcrB*) resistance in environmental *Enterococcus* strains. Notably, none of the *tetL*-positive strains carried all three of *ermB, tcrB*, and *cadA*, suggesting a mutually exclusive distribution pattern and a potentially distinct mechanism of acquisition for *tetL*. These findings reveal novel gene distribution dynamics in environmental *Enterococcus* populations and underscore the importance of integrated surveillance of antimicrobial and metal resistance in human-impacted ecosystems.

## 3. INTRODUCTION

*Enterococcus* species are Gram-positive, facultative anaerobic bacteria that inhabit a wide range of environments, including soil, water, and the gastrointestinal tracts of humans and animals. While many species are commensal, certain strains have emerged as significant opportunistic pathogens due to their ability to acquire and disseminate antibiotic resistance genes. This has led to the increased prevalence of multidrug-resistant (MDR) *Enterococcus* strains, posing major challenges in both clinical and environmental settings.

Aquatic ecosystems are particularly vulnerable to contamination by antibiotic-resistant bacteria due to anthropogenic activities such as wastewater discharge, agricultural runoff, and industrial pollution. The presence of antibiotic-resistant *Enterococcus* strains in water bodies raises concerns about their potential role in the dissemination of resistance genes through horizontal gene transfer. Additionally, heavy metals such as copper and cadmium, commonly found in polluted environments, have been implicated in co-selection for antibiotic resistance due to shared resistance mechanisms.

Given the rising prevalence of MDR *Enterococcus* strains in aquatic environments, this study aims to assess their antibiotic and heavy metal resistance patterns in water bodies of Vladivostok. By analyzing resistance profiles and identifying associated genetic determinants, we seek to enhance our understanding of microbial contamination in water sources and its broader implications for environmental and public health.

## 4. MATERIAL AND METHODS

### 4.1. Sample Collection and Bacterial Isolation

Water samples were collected from two locations subject to strong anthropogenic impact: the estuary of the Vtoraya Rechka River (43.160456° N, 131.905963 ° E) and Zolotoy Rog Bay (43.112913° N, 131.890369° E) in Vladivostok (Figure 1). A total of 20 samples were obtained, yielding 740 bacterial colonies. Among these, 130 morphologically distinct strains were selected for further study. All isolates were confirmed as Gram-positive and catalase-negative. Of the 130 strains, 70 tested positive for the reductase test, suggesting that they might belong to the genus *Enterococcus*.

**Figure 1.**
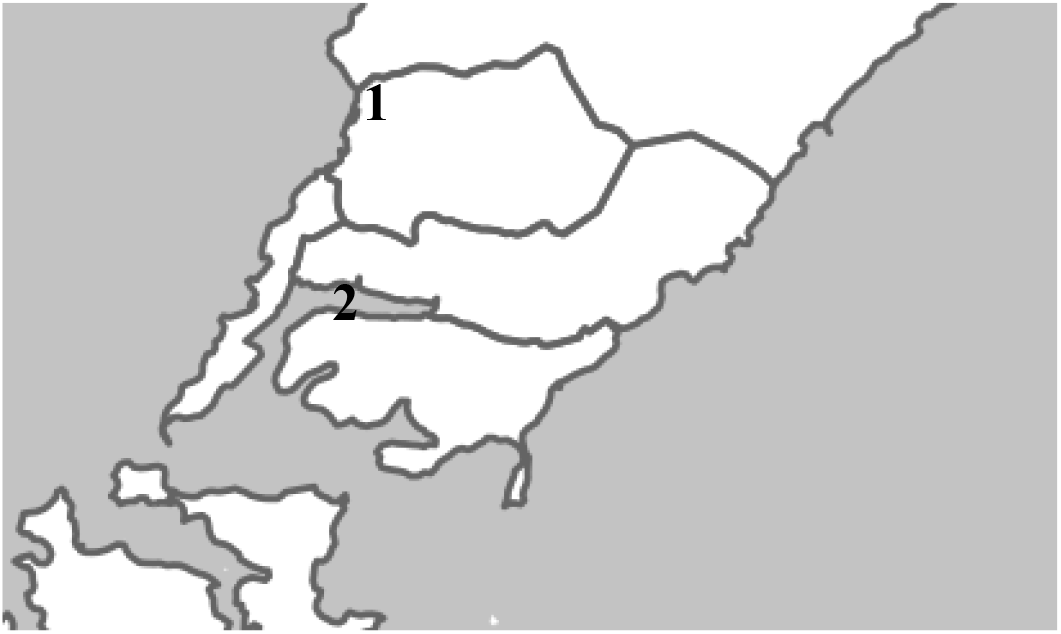
Zolotoy Rog Bay and Vtoraya Rechka River as sample collection sites

### 4.2. PCR and Electrophoresis

Gene-specific PCR primers were used to amplify fragments of the following target genes: 16S1/16S2, *ermB, tetL, tcrB*, and *cadA* (Table 1). Each gene was amplified using tailored thermal cycling conditions adapted from previous studies (Table 2).

**Table 1.**
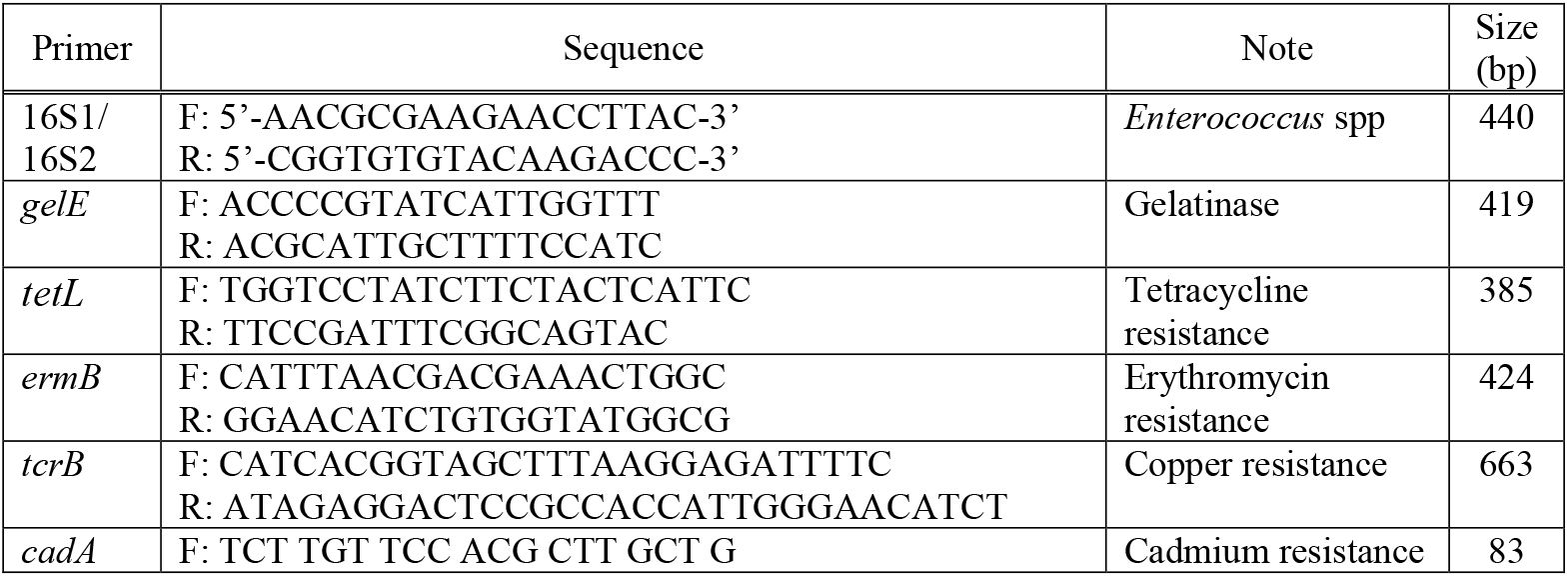
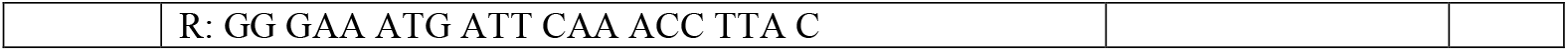
Primers.

**Table 2.** Thermal cycling conditions.

| Primer | Step | Temperature (°C) | Time | Cycle |
| --- | --- | --- | --- | --- |
| 16S1/<br>16S2 | Initial denaturation | 95 | 4 min | 1 |
|  | Denaturation | 95 | 30 sec | 30 |
|  | Annealing | 55 | 1 min | 30 |
|  | Extension | 72 | 1 min | 30 |
|  | Final amplification | 72 | 7 min | 1 |
| *gelE* | Initial denaturation | 92 | 3 min | 1 |
|  | Denaturation | 92 | 5 sec | 5 |
|  | Annealing | 59 | 5 sec | 5 |
|  | Extension | 72 | 5 sec | 5 |
|  | Denaturation | 92 | 1 min | 25 |
|  | Annealing | 59 | 1 min | 25 |
|  | Extension | 72 | 1 min | 25 |
|  | Final amplification | 72 | 10 min | 1 |
| *tetL* | Initial denaturation | 92 | 4 min | 1 |
|  | Denaturation | 92 | 30 sec | 30 |
|  | Annealing | 53 | 1 min | 30 |
|  | Extension | 72 | 1 min | 30 |
|  | Final amplification | 72 | 7 min | 1 |
| *ermB* | Initial denaturation | 94 | 5 min | 30 |
|  | Denaturation | 94 | 1 min | 30 |
|  | Annealing | 51 | 1 min | 30 |
|  | Extension | 72 | 1 min | 30 |
|  | Final amplification | 72 | 5 min | 1 |
| *tcrB* | Initial denaturation | 95 | 5 min | 1 |
|  | Denaturation | 94 | 30 sec | 30 |
|  | Annealing | 55 | 30 sec | 30 |
|  | Extension | 72 | 1 min | 30 |
|  | Final amplification | 72 | 10 min | 1 |
| *cadA* | Initial denaturation | 95 | 5 min | 1 |
|  | Denaturation | 94 | 30 sec | 30 |
|  | Annealing | 55 | 30 sec | 30 |
|  | Extension | 72 | 1 min | 30 |
|  | Final amplification | 72 | 10 min | 1 |

**Table 3.** Electrophoresis results.

| # | Code of strain | 16S RNA | *gelE* | *tetL* | *ermB* | *tcrB* | *cadA* |
| --- | --- | --- | --- | --- | --- | --- | --- |
| 1 | RV1.10.15 | + | + | + | - | - | - |
| 2 | RV1.11.4 | + | - | - | - | + | - |
| 3 | RV1.11.14 | + | + | + | + | - | - |
| 4 | RV1.14.8 | + | - | - | - | + | - |
| 5 | RV1.14.12 | + | + | - | - | - | - |
| 6 | RV1.21.6 | + | - | - | - | + | - |
| 7 | RV1.23.9 | + | - | - | - | + | - |
| 8 | RV1.23.11 | + | - | - | - | - | - |
| 9 | RV1.30.19 | + | + | - | + | - | - |
| 10 | RV1.3.25 | + | + | - | + | + | + |
| 11 | RV1.22.31 | + | + | - | + | + | + |
| 12 | RV1.22.36 | + | - | - | + | + | + |
| 13 | RV1.24.38 | + | + | - | - | - | - |
| 14 | RV1.24.42 | + | - | - | - | - | - |
| 15 | RV1.24.44 | + | + | - | - | - | - |
| 16 | RV2.11.10 | + | + | - | - | - | - |
| 17 | RV2.21.17 | + | - | - | - | - | - |
| 18 | RV2.10.19 | + | + | + | - | - | - |
| 19 | RV2.10.20 | + | - | - | - | + | - |
| 20 | RV2.10.24 | + | + | + | - | + | + |
| 21 | RV2.24.1 | + | - | - | - | - | - |
| 22 | RV2.24.6 | + | + | - | + | - | - |
| 23 | RV2.24.9 | + | + | + | + | - | - |
| 24 | RV2.24.12 | + | - | - | + | - | - |
| 25 | RV2.4.4 | + | + | + | + | - | + |
| 26 | RV2.4.5 | + | + | - | + | + | + |
| 27 | RV2.4.8 | + | - | - | - | + | + |
| 28 | RV2.8.2 | + | + | + | - | - | + |
| 29 | RV2.8.11 | + | + | - | - | + | + |
| 30 | RV2.8.12 | + | - | - | - | - | - |

## 5. RESULTS

### 5.1. Identification of the genus *Enterococcus*

Of the 70 strains tested positive for the reductase test, 30 strains were confirmed as the genus *Enterococcus* by 16S rRNA sequencing. These 30 strains were selected for further genotypic analysis. Species-level identification based on morphological and biochemical characteristics indicated that the majority were *E. faecalis* and *E. faecium*, with a minority of *E. durans* and *E. casseliflavus*.

### 5.2. Genotypic characteristics of virulence and antibiotic-heavy metal resistance

Among the 30 confirmed *Enterococcus* strains, the *gelE* gene, which encodes the virulence factor gelatinase, was detected in 57% (*n* = 17). However, only 13% (*n* = 4) of strains exhibited actual gelatin degradation activity, as determined by the gelatin liquefaction assay. This indicates that not all *gelE*-positive strains demonstrated phenotypic expression of gelatinase, suggesting possible post-transcriptional regulation or gene silencing. The distribution of *gelE* did not appear to correlate with species identity.

The *tetL* gene, associated with tetracycline resistance, was detected in 23% (*n* = 7) of the strains. The *ermB* gene, which confers resistance to macrolides such as erythromycin, was detected in 33% (*n* = 10). The *tcrB* gene (copper resistance) was identified in 40% (*n* = 12) of strains. The *cadA* gene (cadmium resistance) was detected in 27% (*n* = 9) (Figure 2).

**Figure 2.**
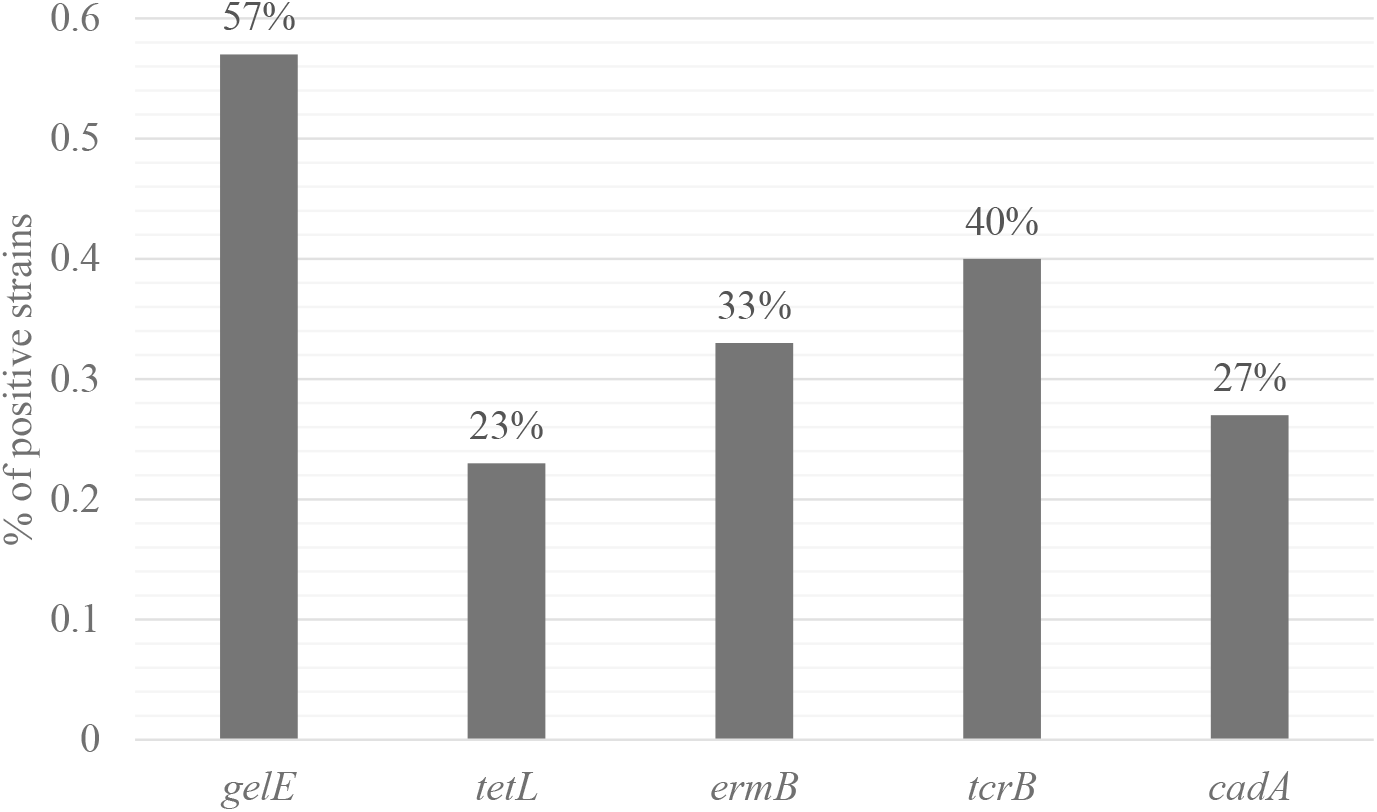
Prevalence of the *gelE, tetL, ermB, tcrB*, and *cadA* genes

### 5.3. Co-occurrence of genes responsible for resistance to antibiotics and heavy metals

Out of the 30 *Enterococcus* strains, 76.7% (*n* = 23) tested positive for at least one resistance gene related to antibiotics or heavy metals. Among these, 46.7% (*n* = 14) carried more than one resistance gene.

Notably, 13.3% (*n* = 4) of strains carried both the *ermB* and *tcrB* genes, suggesting a potential genetic association between copper resistance and macrolide resistance (Figure 3). In addition, a pattern was observed in which strains harboring multiple resistance genes tended to carry a combination of *ermB, tcrB*, and *cadA*. In contrast, the *tetL* gene was frequently absent from these multiresistant strains.

**Figure 3.**
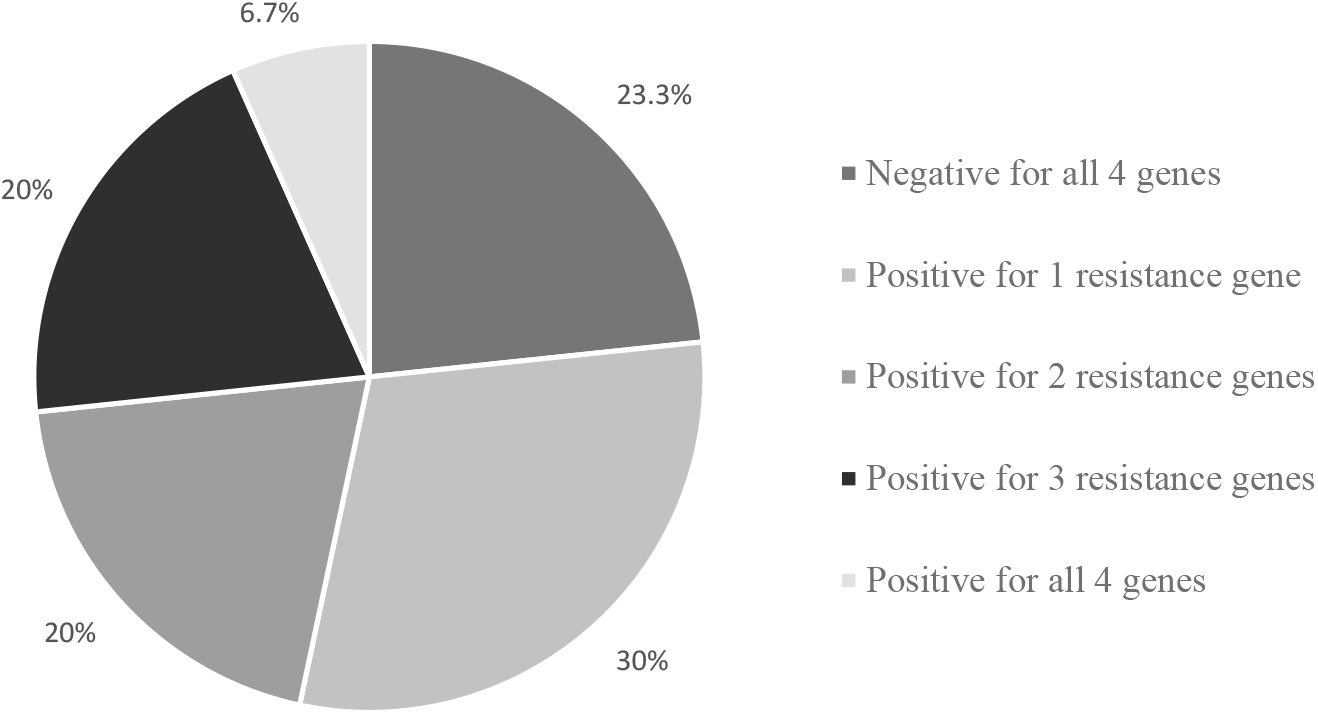
Co-occurrence of the *tetL, ermB, tcrB*, and *cadA* genes

Among the *tetL*-positive strains (*n* = 7), most were associated with no additional resistance genes (*n* = 2), or with only one (*n* = 3) or two (*n* = 2) resistance genes. Importantly, none of the *tetL*-positive strains carried all three of *ermB, tcrB*, and *cadA*, which may indicate a mutually exclusive distribution pattern between *tetL* and the co-occurrence of other resistance genes (Figure 4). This suggests that *tetL* may be acquired or maintained independently, while the other resistance determinants may be co-selected or genetically linked.

**Figure 4.**
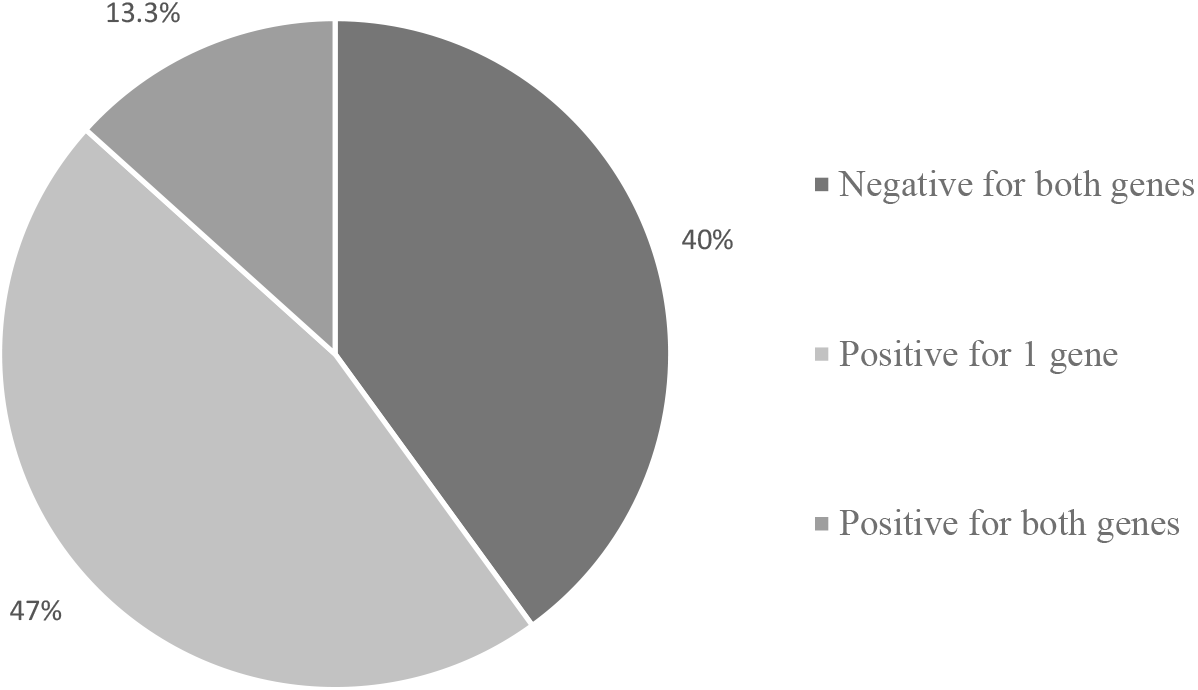
Co-occurrence of the *ermB* and *tcrB* genes

**Figure 5.**
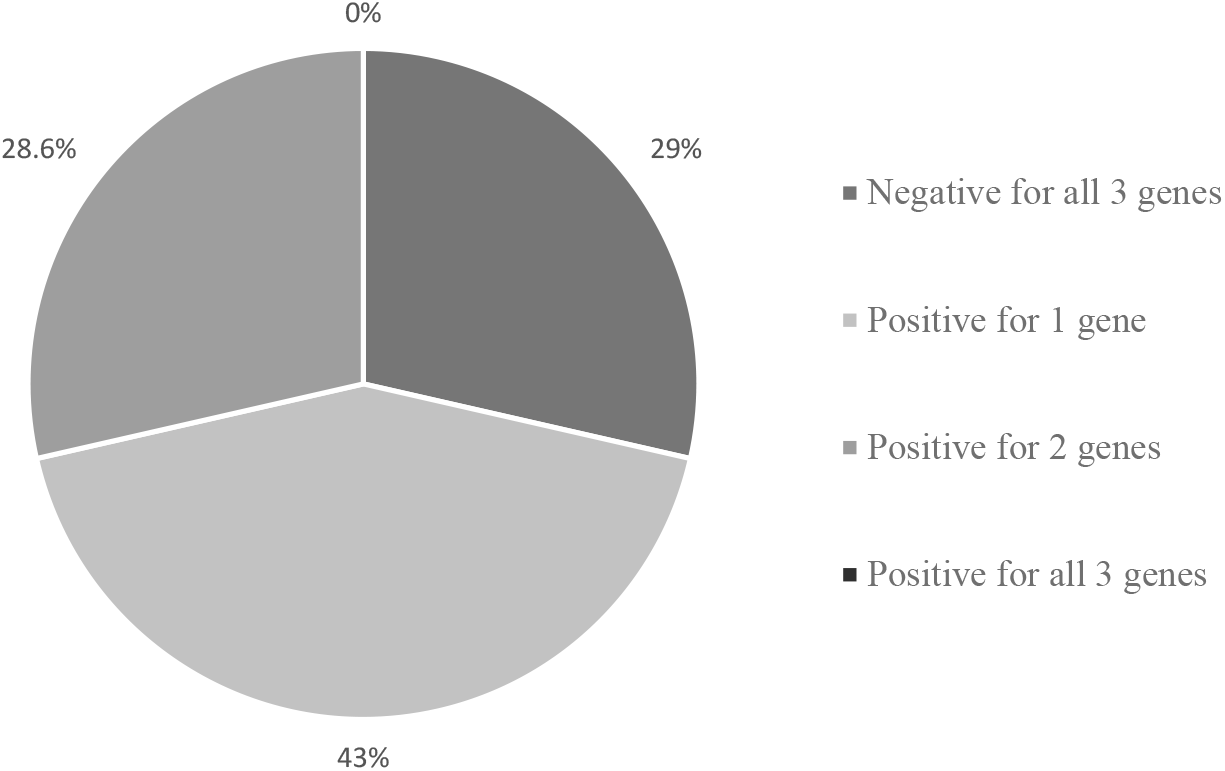
Co-occurrence of the *ermB, tcrB*, and *cadA* genes in *tetL*-positive strains

## 6. DISCUSSION

Genomic analyses of *Enterococcus* strains isolated from Vladivostok’s water bodies revealed that the resistance genes *tetL, ermB, tcrB*, and *cadA*—conferring resistance to tetracycline, erythromycin, copper, and cadmium, respectively.

Notably, the co-occurrence of the *ermB* and *tcrB* genes in 13.3% of strains suggests a potential association between antibiotic and heavy metal resistance.

While many strains demonstrated multigene resistance—most commonly involving *ermB, tcrB*, and *cadA*—*tetL*-positive strains tended to appear in genetic isolation. Specifically, none of the *tetL*-positive strains exhibited simultaneous presence of *ermB, tcrB*, and *cadA*, which suggests a mutually exclusive relationship between *tetL* and this multiresistance pattern. This may imply that *tetL* is acquired or maintained independently of other resistance determinants, while *ermB, tcrB*, and *cadA* may be part of a co-selected or physically linked resistance cluster. This distribution pattern reinforces the importance of considering gene linkage and selection dynamics when assessing resistance dissemination in environmental *Enterococcus* populations.

The intrinsic and acquired resistance mechanisms of *Enterococcus* species enable them to withstand multiple classes of antibiotics, including β-lactams, aminoglycosides, and glycopeptides, which complicate infection management. The widespread resistance, particularly to aminoglycosides such as streptomycin, raises concerns about the possible emergence of vancomycin-resistant enterococci (VRE) in the future.

Future investigations should focus on comprehensive genetic characterization and antibiotic resistance profiling to elucidate the pathogenic potential and environmental adaptability of these strains, thereby informing strategies to mitigate their impact on public health.

## 7. ACKNOWLEDGMENTS

We would like to thank our colleagues and the laboratory staff at Far Eastern Federal University Institute of the World Ocean for their invaluable assistance with sample collection and experimental procedures. We also appreciate the support provided by Sintol (Moscow, Russia) for synthesizing the PCR primers.

